# Protective geometry and reproductive anatomy as candidate determinants of clutch size variation in pentatomid bugs

**DOI:** 10.1101/2022.05.16.492197

**Authors:** Paul K. Abram, Eric Guerra-Grenier, Jacques Brodeur, Clarissa Capko, Michely Ferreira Santos Aquino, Elizabeth H. Beers, Maria Carolina Blassioli-Moraes, Miguel Borges, M. Fernanda Cingolani, Antonino Cusumano, Patrick De Clercq, Celina A. Fernandez, Tara D. Gariepy, Tim Haye, Kim Hoelmer, Raul Alberto Laumann, Marcela Lietti, J.E. McPherson, Eduardo Punschke, Thomas E. Saunders, Jin Ping Zhang, Ian C.W. Hardy

## Abstract

Many animals lay their eggs in clusters. Eggs on the periphery of clusters can be at higher risk of mortality. We asked whether the most commonly occurring clutch sizes in pentatomid bugs could result from geometrical arrangements that maximize the proportion of eggs in the cluster’s interior. Although the most common clutch sizes do not correspond with geometric optimality, stink bugs do tend to lay clusters of eggs in shapes that protect increasing proportions of their offspring as clutch sizes increase. We also considered whether ovariole number, an aspect of reproductive anatomy that may be a fixed trait across many pentatomids, could explain observed distributions of clutch sizes. The most common clutch sizes across many species correspond with multiples of ovariole number. However, there are species with the same number of ovarioles that lay clutches of widely varying size, among which multiples of ovariole number are not over-represented. In pentatomid bugs, reproductive anatomy appears to be more important than egg mass geometry in determining clutch size uniformity. In addition, within this group of animals that has lost most of its variation in ovariole number, clutches with a broad range of shapes and sizes may still be laid.

## Introduction

A fundamental aspect of the life history of animals is how many offspring they produce per bout of reproduction (Godfray et al. 1991). Some egg-laying animals, most notably insects, fix their eggs to substrates in two- or three-dimensional clusters of multiple eggs. Several hypotheses have been put forward to explain the benefit of laying eggs in clusters for both mothers and offspring (reviewed in Faraji et al. 2002; see also: Courtney 1984; Friedlander 1985; Clark and Faeth 1998; Gamberale and Tullberg 1998). For mothers, clustering eggs in particularly suitable locations instead of laying them individually could reduce costs (e.g., time, energy, predation/infection risk) of finding suitable oviposition sites. For individual offspring, benefits of developing in a cluster of eggs can include protection against desiccation, improved thermoregulation, Allee effects of aggregated feeding after emergence, improved efficacy of aposematism by aggregated offspring after emergence and/or saturating predator consumptive ability (Stamp 1980). While these factors may help explain why eggs are clustered rather than laid singly and, to some degree, why some species typically produce larger clutches than others, they do not necessarily address the related questions of how many eggs should be laid per egg cluster, what the shape the egg clusters should be and how consistent clutch size should be within species. These questions are interrelated as, for instance, the shape of an egg cluster is to an extent constrained by the number of eggs comprising the cluster.

There is evidence that the spatial arrangement of eggs within clusters can influence offspring survival, with individuals on the margins having lower survival, in a manner similar to ‘selfish herd’ effects observed across many taxa of both sessile and motile organisms that form aggregations (Hamilton 1971; McDowall and Lynch 2019). In insects, eggs on the insides of clusters may be less susceptible to predation and parasitism (Weseloh 1972; Friedlander 1985; Gross 1993; Hondo et al. 1995; Mappes and Kaitala 1994; Mappes et al. 1997; Kudo 2001; Kudo 2006; Deas and Hunter 2012) and may also suffer lower mortality due to desiccation (Clark and Faeth 1998). Intriguingly, some species of true bugs (Insecta: Hemiptera) lay smaller eggs on the periphery of egg masses – where they suffer higher rates of predation – than on the inside, presumably reflecting a reduced maternal investment in eggs that are more vulnerable (Mappes and Kaitala 1994; Mappes et al. 1997; McLain and Mallard 1991; Kudo 2001; Kudo 2006). There is a further implication of these findings: because the proportion of eggs on the periphery of a cluster depends on both the number of eggs in the cluster and the shape in which the cluster is arranged, there may be optimal clutch sizes and shapes that minimize the mean per-egg vulnerability by maximizing the proportion of eggs that are ‘protected’ in the interior of a cluster.

For objects such as eggs that are approximately circular in a two-dimensional plane, the dense-packing arrangement that minimizes exposed surface area at the periphery is the hexagonal lattice pattern (Chang and Wang 2010)^1^. Further, complete hexagonal arrangements of circular eggs are only possible for certain numbers of eggs (Figure 1). Thus, when eggs are arranged in the closest possible approximation of a hexagon, there are certain clutch sizes for which adding one more egg would increase the proportion of eggs that are protected within the interior of the egg mass (hereafter, “protection efficiency”) and other clutch sizes for which adding an egg would decrease the protection efficiency. This implies that for hexagonally arranged clusters of eggs, there are ‘adaptive peaks’ of clutch size where protection efficiency could be locally maximized (Figure 1). These peaks are contained in the following integer sequence *a*(*n*), where *n* is the position in the sequence:

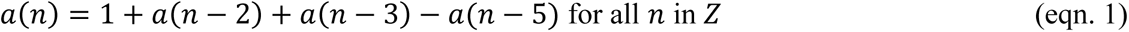

**Figure 1.**
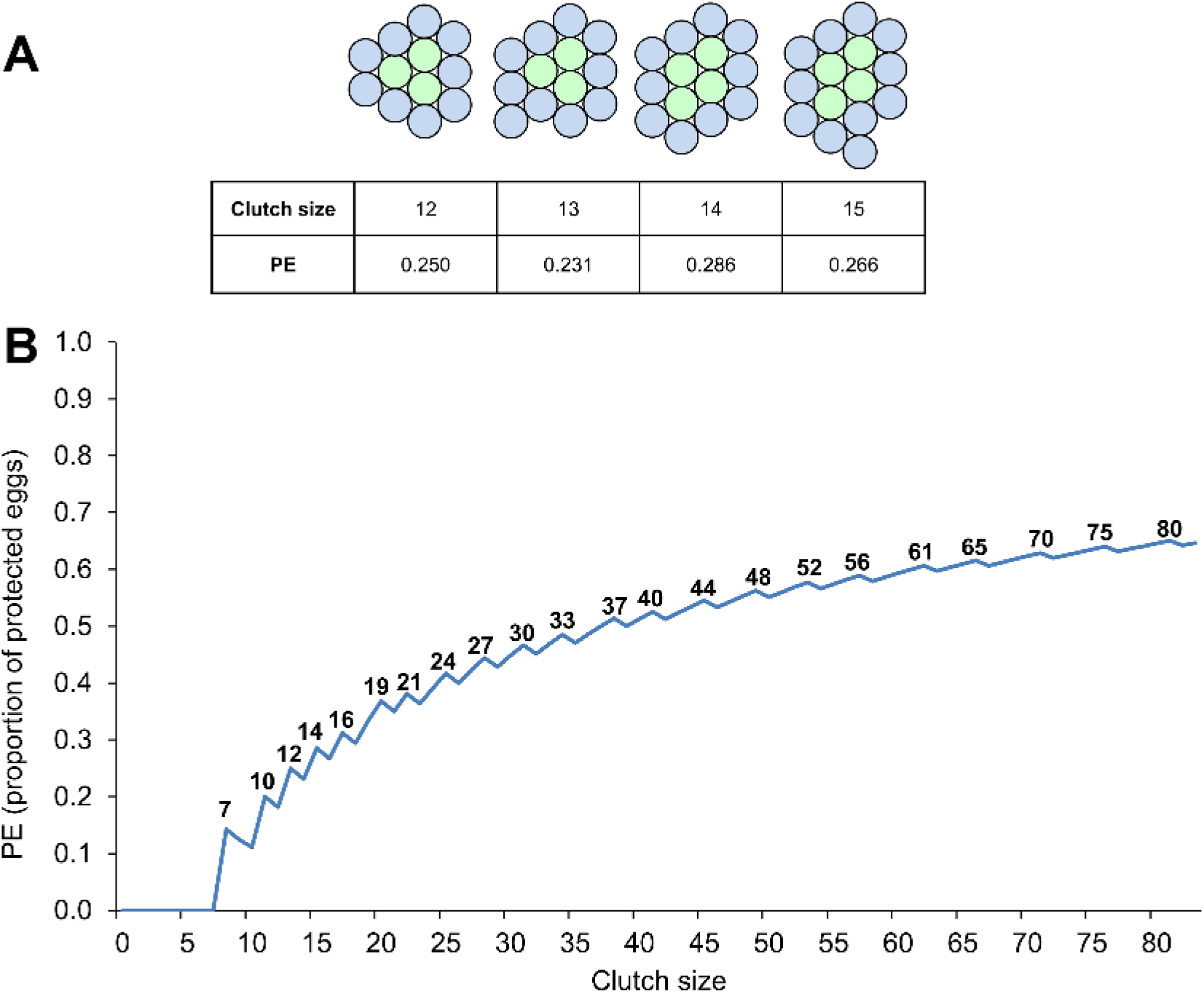
How protection efficiency (PE), the number of eggs protected on the inside of a clutch, varies with clutch size for shapes most closely approximating a hexagon. (A) Illustration for selected clutch sizes. Adding eggs to the outside of an egg mass increases the number of protected eggs (shown in green), and PE, when increasing clutch size from 13–14, but PE declines for clutch size increases of 12–13 or 14–15 because the total number of eggs increases without increasing the number protected on the inside. (B) The “Sawtooth hypothesis” showing PE for all clutch sizes up to 82. Local “peaks” in PE are labeled with the corresponding clutch size.

*a*(*n*) is a known integer sequence describing the number of partitions of *n* (i.e., a multiset of positive integers that sum to *n*) into at most three parts; where *n* is the position in the integer sequence and *Z* is length of the integer sequence (OEIS 2022). If these peaks in protection efficiency are indeed adaptive, geometry could be a determinant of clutch size evolution, with natural selection leading to clutch sizes occurring disproportionally frequently at these peaks. In addition, because the proportional increase in protection efficiency is greater for smaller clutch sizes (Figure 1), uniform clutch size might be expected to evolve more often when clutches are smaller. Finally, the strength of selection towards these peaks (and thus the uniformity of clutch size) could vary interspecifically, for instance, depending on the strength of predation/parasitism pressure on eggs of a particular species and how focused it is on eggs on the periphery of clusters.

Stink bugs (Hemiptera: Pentatomidae) provide a spectacular example of insects that lay clustered egg masses (Figure 2). Stink bugs are distributed globally across a wide range of ecosystems and have a wide array of life-histories, including herbivorous and predatory species (McPherson and McPherson 2000; Panizzi et al. 2000; Grazia et al. 2015). While conducting research on a thoroughly studied pest insect, the brown marmorated stink bug, *Halyomorpha halys* (Hemiptera: Pentatomidae), it was noticed that most egg masses consisted of 28 eggs (PKA, EGG, JB; pers. obs.), as observed in past studies of this species (e.g., Nielsen et al. 2008). However, another pentatomid species being studied, *Podisus maculiventris*, laid clutches with highly variable numbers of eggs (Torres-Campos et al. 2016). Such divergent patterns of clutch size uniformity or variability have also been noted by other stink bug researchers (e.g., McPherson 1976; Panizzi and Herzog 1984; De Clercq and Degheele 1997; Matesco et al. 2009), but the factors responsible for these patterns are not well understood.

**Figure 2.**
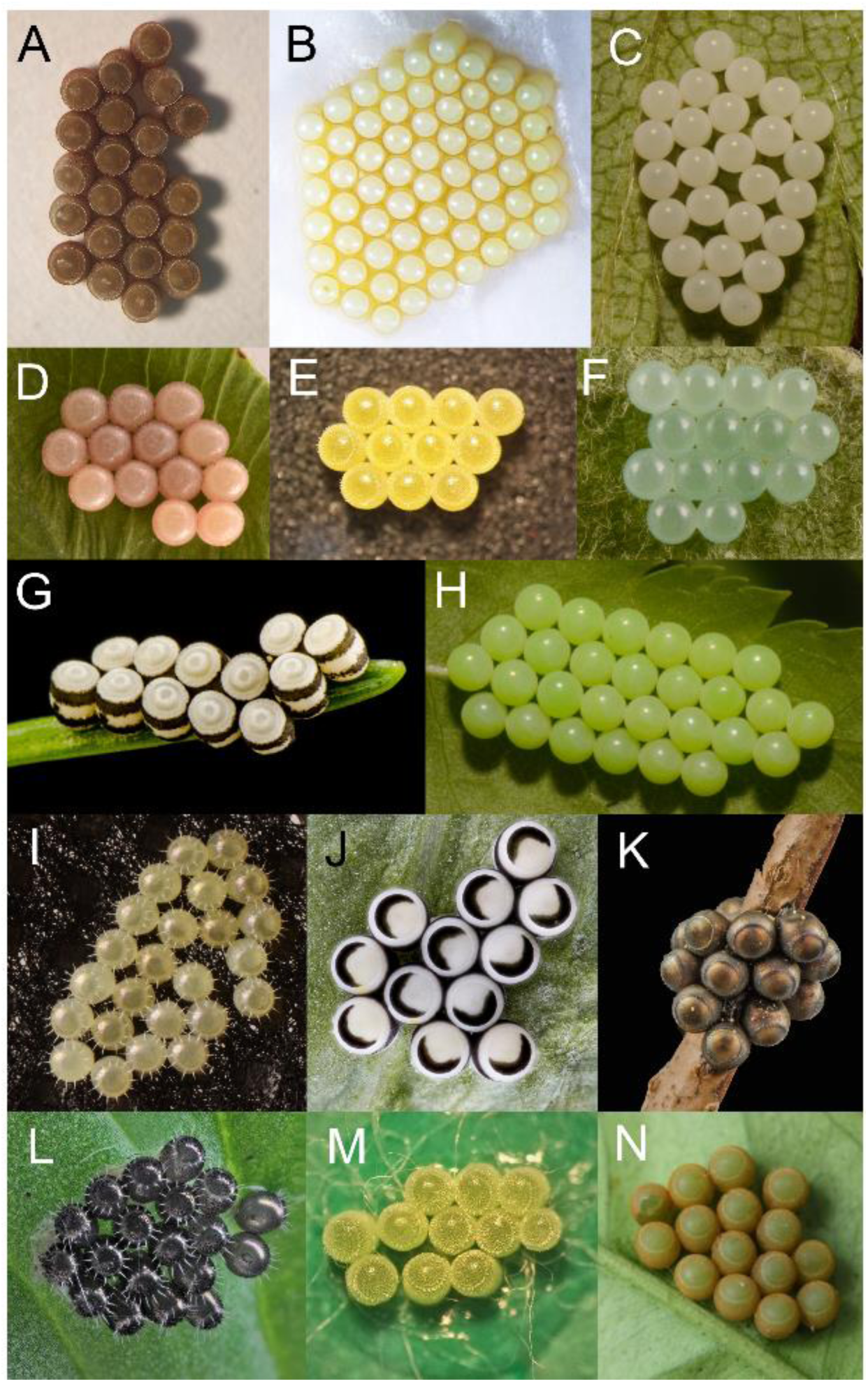
Examples of stink bug egg mass geometry: (A) *Chinavia impicticornis*; (B) *Nezara viridula*; (C) *Halyomorpha halys*; (D) *Dolycoris baccarum*; (E) *Euschistus heros*; (F) *Pentatoma rufipes*; (G) *Chlorochroa pinicola*; (H) *Palomena prasina*; (I) *Arma custos*; (J) *Murgantia histrionica*; (K) *Rhaphigaster nebulosa*; (L) *Podisus maculiventris*; (M) *Dichelops melacanthus*; (N) *Glaucias amyoti*. All photographs were taken by the authors except for three that were sourced from Wikimedia Commons: (J) Public domain; (L) Rsbernard, CC BY-SA 4.0 and (N) Groovybandit, CC BY-SA 4.0.

One possible constraining factor on clutch size variability is reproductive anatomy. Most female stink bugs have seven ovarioles per ovary, giving a total of 14 (Table S1 and references therein). Each ovariole can contain one or more oocytes maturing at a given time. Ovariole numbers have been hypothesized to be determinants of clutch sizes in pentatomids (Kiritani and Hokyo 1965; Panizzi and Herzog 1984; Matesco et al. 2008; Matesco et al. 2009) as well as total lifetime reproductive output in other insect lineages (e.g., Price 1974; Ware et al. 2008; Sarikaya et al. 2019; Church et al. 2021). Church et al. (2021) recently reported that several insect taxa, including the Pentatomidae, have evolved invariant, or near-invariant, ovariole numbers. How near-invariant ovariole numbers might influence clutch size variation is currently unclear.

Following from the arguments outlined above regarding egg mass geometry, protection efficiency and reproductive anatomy, we generated two different – but not mutually exclusive – hypotheses to explain variation in stink bug clutch sizes. First, the most frequent clutch sizes (within and among species) will be determined by geometric optimality for protection efficiency and will fall on adaptive peaks predicted in Figure 1 (hereafter, the “sawtooth hypothesis”, named after the serrated form of the relationship). According to this hypothesis, clutches should fall more often on adaptive peaks when clutch sizes are small (above a critical minimum needed to protect one egg), with the arrangements of their component eggs tending towards optimal packing. Second, the most frequent clutch sizes (within and among species) are determined by a constraint imposed by ovariole number and will therefore be equal to multiples of that number (hereafter, the “ovariole multiple hypothesis”).

To characterize variation in clutch parameters in the Pentatomidae and test both hypotheses, we collected clutch size data from research groups studying stink bugs in several countries and also examined photographs of stink bug egg masses from online searches, publications, and laboratory cultures. Our study may be the first to consider whether and how the geometry of egg clusters and reproductive anatomy could influence the evolution of animal clutch size and its uniformity.

## Materials and Methods

The ‘sawtooth’ and the ‘ovariole multiple’ hypotheses were evaluated by analyses of two different datasets: the first evaluated clutch shape; the second focused on clutch size.

Unless otherwise indicated, all hypothesis testing analyses were carried out in R statistical software (R Core Team 2019).

### Part 1: Clutch shape variation

First, we tested whether the shape of stink bug egg masses conformed to predictions of the sawtooth hypothesis, and whether egg masses with clutch sizes predicted by the sawtooth hypothesis had higher indices of protective geometry than those predicted by the ovariole multiple hypothesis.

#### Data collection

We extracted data from photographic images of stink bug egg masses collected from online searches, publications, and our own laboratory cultures. For web searches, we entered “Species name + eggs” as keywords in Google Images and inspected the results. To be included in the database, a photograph had to comply with all of the following criteria: (i) have been taken from directly above the clutch such that non-overlapping eggs did not appear to overlap; (ii) have high enough resolution such that individual eggs were clearly identifiable (not unduly pixelated); (iii) clutches had been laid on a relatively flat surface; and (iv) the contours of hatched eggs were not obscured by opercula or nymphs. We included species for which we were able to obtain at least three egg mass images; for species with more photographs available, we sampled up to five randomly selected photographs so that the per-species replication would be relatively balanced (3-5 egg masses per species). The final analysis included 78 egg masses from 19 pentatomid species in 15 genera.

#### Determination of ovariole numbers

Using the ovariole number database of Church et al. (2021) as a starting point, we conducted a literature search of ovariole numbers per ovary in the Pentatomidae. While ovariole number per ovary from most pentatomid species in the literature was seven, in agreement with all entries for pentatomids in the Church et al. (2021) database, we found that at least two species in the tribe Strachinii only have six ovarioles per ovary (Benedek 1968, Grodowitz et al. 2019). Follow-up dissections of two additional species by the current study’s co-authors also found that *Murgantia histrionica*, which is in the tribe Strachinii, also has six ovarioles per ovary (A. Cusumano, personal observations). Dissections of *Cappaea tibialis* to determine ovariole number were also done *a posteriori* (after inspecting the data); these found that *C. tibialis* has eight ovarioles per ovary (J. Zhang, personal observations). Thus, in subsequent analyses, ovariole number for all species was set to seven as a default, with the exception of *C. tibialis* and species in the tribe Strachinii, for which ovariole number was set to six and to eight, respectively.

#### Image analysis

Clutch images were analyzed individually using ImageJ v1.53c. Eggs were first delimited using the ‘oval selection’ tool and recorded individually as regions of interest (ROI). Using each egg ROI, we calculated the average egg diameter. By combining all egg ROIs, we measured the total area occupied by eggs within each clutch. We then traced the clutch perimeter to measure both total perimeter and total clutch area (i.e., area occupied by eggs and the gaps between them). Subsequently, we used a custom macro to identify and mark the centroid of each egg. These marks then allowed us to draw a line between neighboring eggs and later measure the average distance between egg pairs. We considered two eggs as neighbors if a line could be drawn between their centers without intersecting the perimeter of a third egg. Following this method, we extracted three main variables: protection efficiency (PE), egg aggregation, and clutch circularity. See Figure S1 for an example of ROIs drawn on an egg mass photograph in ImageJ.

Protection efficiency was measured as the proportion of eggs that were on the inside of the clutch (i.e., eggs that did not intersect with the clutch perimeter). Observed protection efficiency values were compared to theoretical (geometric) optima at the corresponding clutch sizes to measure the protection differential (PD = Observed PE – Optimal PE). A differential of 0 indicates that the observed protection efficiency is optimal under the sawtooth hypothesis.

Egg aggregation, a measure of how tightly packed eggs were, was calculated as an index by dividing the average egg diameter by the average distance between neighbouring eggs. Theoretically, the shortest possible distance between the centres of two (equally sized) circles is equal to their diameter since the distance of their two centers would equal the sum of their radii. Therefore, two eggs touching each other would have a maximal aggregation value of 1 (although we did measure values slightly higher than 1 when two eggs naturally overlapped or were deformed when pressed against one another).

Clutch circularity was measured using the equation embedded in ImageJ (4 × Pi × (Area **÷** Perimeter^2^). Values closer to 1 indicate shapes that tend towards perfect circles and, thus, towards an optimal Area:Perimeter ratio.

#### Quantifying taxonomic relatedness among species

To account for phylogenetic non-independence resulting from the shared ancestry of the species in our dataset, we first needed to establish estimates of the phylogenetic relatedness among species. A recent study based on several mitochondrial and nuclear genes suggests that the family Pentatomidae requires comprehensive taxonomic revision (Roca-Cusachs et al. 2022), but at present there is no consensus phylogeny that includes most of the species in our dataset. In the absence of an applicable consensus phylogeny for the Pentatomidae, and recognizing that taxonomic relationships are likely to be revised in the future, we provisionally used taxonomic relationships as a proxy for phylogenetic relatedness (e.g., Straub et al. 2011). We constructed a distance matrix based on taxonomic distances among each species pair and assumed equal branch lengths (=1) for each taxonomic division (subfamily, tribe, genus, species) (Straub et al. 2011; Abram et al. 2021) with each successive division represented as a polytomy. We assigned a single, ‘placeholder’ tribe designation for genera in the subfamily Asopinae and for the genus *Thyanta* which has not been placed in tribes.

#### Clutch shape analyses

First, we plotted the protection efficiency of the 78 stink bug egg masses against optimal protection efficiencies predicted by the sawtooth hypothesis and qualitatively examined their correspondence. Then, to examine whether protection differential changed with clutch size (i.e., if larger clutches have proportionally better-protected eggs), accounting for shared ancestry (see “Quantifying taxonomic relatedness among species”), we fitted a Phylogenetic Least Squares (PGLS) model (Freckleton et al. 2002) using the pgls() function in the “caper” package (Orme et al. 2012), with protection efficiency as the response variable and log-transformed clutch size (to meet the parametric assumptions of the analysis) as the independent variable.

Next, we examined whether clutches of a size that could lie on an ‘adaptive peak’ predicted by the sawtooth hypothesis actually had better indices of protective geometry than clutch sizes predicted by the ovariole multiple hypothesis, neither of the two hypotheses, or both hypotheses. Clutch sizes conforming to the sawtooth hypotheses were those that were contained in the integer sequence outlined in eqn. 1. Clutch sizes conforming to the ovariole multiple hypothesis were those that were multiples of each species’ per ovary ovariole number. We used PGLS models to test whether protection differential, circularity, or aggregation index varied among egg masses with sizes that corresponded to each hypothesis, neither or both.

#### Intraspecific variation in clutch shape

To examine more detailed patterns of intraspecific variation in to what degree protection efficiency of egg masses conformed to the sawtooth hypothesis, we analyzed additional photographs of at least 18 and up to 50 egg masses produced in laboratory cultures of five species that were available and producing egg masses at the time we initiated the analysis in 2021: *H. halys* (n = 24; British Columbia, Canada), *Chinavia impicticornis* (n = 18; Brazil), *Euschistus heros* (n = 25; Brazil), *Nezara viridula* (n = 50; New Zealand and Italy), and *Cuspicona simplex* (n = 50; New Zealand). Protection efficiency was measured as described above, and we qualitatively evaluated resulting fits against the sawtooth hypothesis. We chose three egg masses per species that had the same clutch size and manually produced vector drawings of each to qualitatively examine how individual species can lay the same clutch sizes in different conformations.

### Part 2: Clutch size variation

We next tested whether the clutch sizes and degree of clutch uniformity of stink bugs conformed better to predictions of the sawtooth hypothesis or the ovariole multiple hypothesis. Because clutch size data were more readily available than egg mass images, this dataset was much larger (more species and more egg mass data per species) and more comprehensive than the photographic dataset.

#### Data collection

We collected data on the number of eggs in a total of 21,002 clutches, representing 51 stink bug species, from 13 research groups conducting collections in seven countries in North America (Canada, USA), South America (Brazil, Argentina), Europe (Belgium, Italy, Switzerland), Oceania (New Zealand), and Asia (China, Japan). One dataset on *P. maculiventris* was excluded at the outset because it was an order of magnitude larger than any others and contained extreme clutch values that could not be verified. With one exception (*N. viridula,* studied in Belgium), data were not initially collected for this study but derived from several published and currently unpublished studies of pentatomid biology, carried out by the co-authors of the current article, in which clutch size information was recorded. Of the 88 remaining datasets, 10 were records of naturally occurring egg masses in the field and 78 were from egg masses collected from laboratory cultures reared under a range of conditions (typically, 16:8 L:D photoperiod, 20-25°C). From the initial 51 species, five were excluded from further consideration due to small sample size (< 30 observations of clutch size). This resulted in a total of 20,906 egg masses in the analyzed dataset. There were one to seven independently collected datasets for each of the 46 species considered, with at least two datasets for 13 species. For all subsequent analyses, we pooled multiple datasets for each species, accepting that the range of variation recorded for each species could be affected by sampling effort and the range of conditions under which data were collected. However, patterns of clutch size variation and uniformity were generally similar among datasets (including laboratory *vs.* field studies) for the same species (Figure S2), with the exception of species that laid large and highly variable-sized egg masses, which did show some between-dataset heterogeneity (e.g., *N. viridula*). We note that the overall set of data that we explored is unlikely to be a fully random sample of clutch size variation within the Pentatomidae; the species that are studied by researchers, and that have been contributed to the current study, are typically common, economically significant pests (or their close relatives) and amenable to laboratory rearing.

For each of the 46 species, we calculated the mean and modal clutch size. We also calculated for each species a percentage index that we term ‘clutch size uniformity’: the number of clutches with the modal clutch size divided by the total number of clutches observed, multiplied by 100.

#### Clutch size analysis

We tested whether clutch sizes observed in each species conformed to predictions arising from our two hypotheses, here formulated in terms of clutch size: (i) Sawtooth hypothesis: clutch sizes that optimize geometric protection efficiency are more common than would be expected if clutch sizes are Poisson-distributed (i.e. randomly – the null scenario) around the observed mean; (ii) Ovariole multiple hypothesis: clutch sizes that are multiples of a species’ ovariole number per ovary are more common than would be expected if clutch sizes are Poisson-distributed around the observed mean. Each clutch was designated as conforming to each of the two hypotheses as described above (see “Clutch shape analyses”).

For tests of both hypotheses for each species, we then calculated the expected number of clutches that would fall on sawtooth or ovariole multiple clutch sizes, over the observed range of clutch size under the null scenario. 2 × 2 Chi-squared tests of independence were then performed for each hypothesis and each species to test whether the observed distribution of clutch sizes differed significantly from the null (Poisson) scenario. Using the spreadsheet tool developed by McDonald (2014), we used the Benjamini-Hochberg procedure (Benjamini and Hochberg, 1995) to control for multiple comparisons, considering the group of tests for each of the two hypotheses as separate families of tests. The family-wide α value (i.e., the false discovery rate) was set at 0.10. To graphically present how over- or under-represented sawtooth and ovariole-multiple-conforming clutch sizes are for each species, we calculated their “Relative representation”, measured as a percentage, equal to the observed number of clutches fitting the hypothesis, minus the number that would be expected under the null (Poisson) scenario, divided by the total number of clutches in the dataset.

#### Clutch uniformity analysis

To test the prediction of the sawtooth hypothesis that more uniform clutch sizes should occur more often for species with smaller clutches, we treated clutch size uniformity as the response variable and mean clutch size as the explanatory variable. Both variables were log-transformed to meet assumptions of normality and homoscedasticity. Using a taxonomic distance matrix, generated for the clutch size dataset as described above (see “Clutch shape analysis”), we then fitted a PGLS model with clutch size uniformity as the response variable and log-transformed clutch size as the independent variable. An additional quadratic term (log [mean clutch size]^2^) was added to the model after we detected non-linearity in the residuals plot of the original model. Inspection of residual and qq-plots, as well as comparisons of Akaike’s Information Criterion between the model containing only the linear term (AIC=108.57) compared to that with the additional quadratic term (AIC=103.74) confirmed that the relationship was most parsimoniously described by the model containing the quadratic term (significance of quadratic term: p = 0.012).

#### Taxonomic patterns of clutch size and uniformity

Using the taxonomic distance matrix, we calculated Pagel’s λ (Pagel 1999), which is an index of phylogenetic signal (here, applied to a phylogeny based on taxonomic relationships), to characterize the extent to which closely related pentatomid species have similar clutch size means, uniformity, and relative representation under each of our two hypotheses. Pagel’s λ generally varies between 0 and close to 1, where 0 indicates no phylogenetic signal (traits evolve independent of phylogeny) and a value close to 1 indicates that traits co-vary as would be expected under an assumption of a pure Brownian motion model of evolution, where the phylogenetic relationships among species determine how their clutch size traits co-vary. Pagel’s λ performs well relative to other indices of phylogenetic signal when phylogenies are unresolved or there is suboptimal branch length information available (Münkemüller et al. 2012; Molina-Venegas and Rodríguez 2017), as is the case in our study. To estimate Pagel’s λ, we used the “phylosig” function in the “phytools” package. Likelihood ratio tests were used to test for taxonomic signals for each trait, i.e. whether λ deviated significantly from zero.

## Results

### Part 1: Clutch shape variation

#### Clutch shape analyses

The protection efficiency of only four out of 78 egg masses, laid by three different species (*Dichelops furcatus* - 1, *Euschistus conspersus* - 2, *Palomena prasina* - 1), were situated on the peaks of geometric optimality predicted by the sawtooth hypothesis (Figure 3A). In addition, the observed protection efficiency of egg masses was suboptimal over the entire range of observed clutch sizes (Figure 3A), with egg masses protecting, on average, 10.7% fewer eggs than predicted if egg masses were to be laid in such a way as to optimize protection. However, as clutch size increased, the protection efficiency of egg masses was closer to optimal values (Figure 3B).

**Figure 3.**
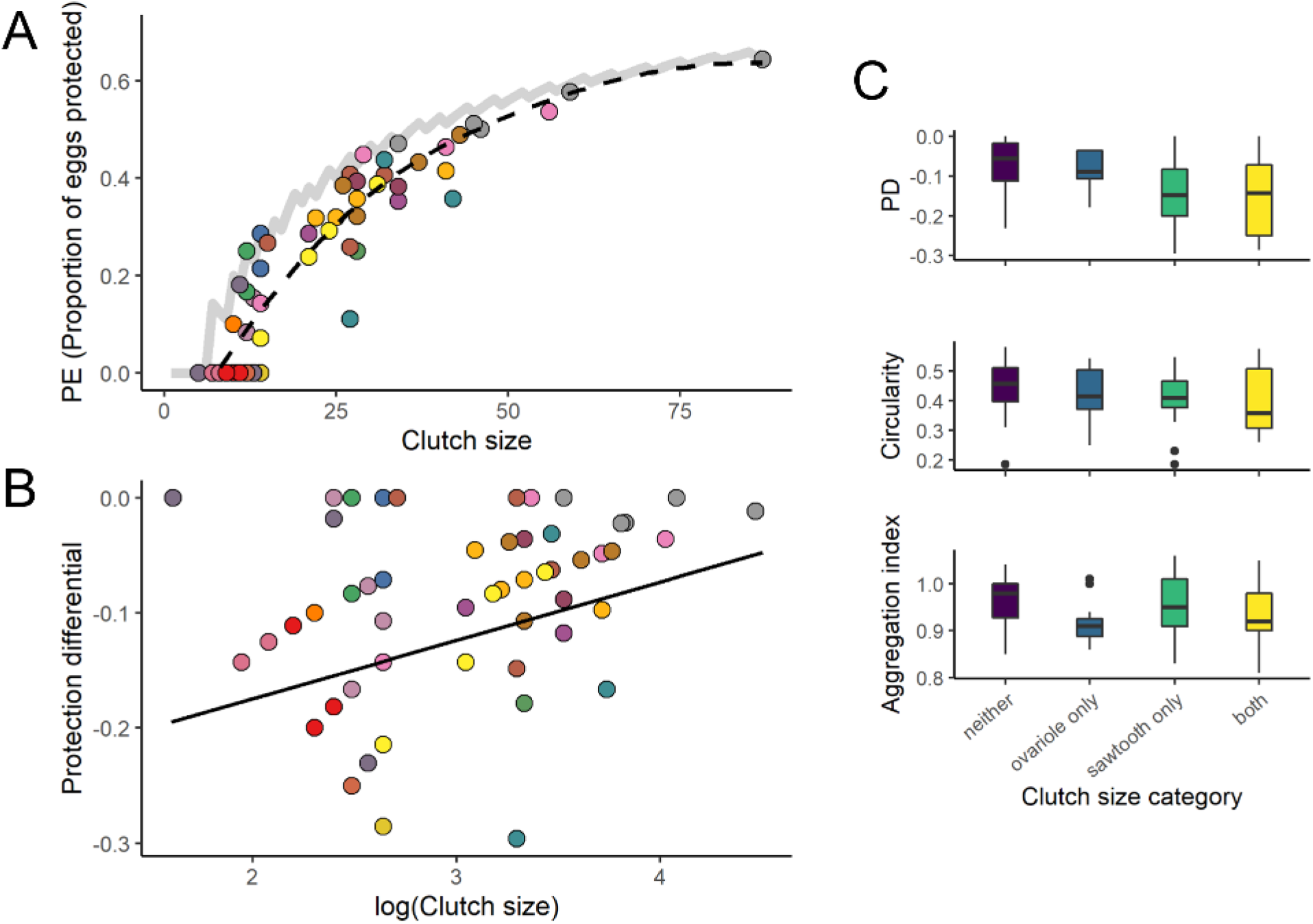
The arrangement of 78 pentatomid egg masses (from 19 species) in relation to the sawtooth and ovariole multiple hypotheses. (A) The protection efficiency (PE), the proportion of eggs protected in the interior of the egg mass, compared to optimal PEs predicted by the sawtooth hypothesis (grey line); the dotted black line is a local polynomial regression (loess) curve (smoothing parameter, α = 1). (B) The relationship between protection differential (actual PE of egg mass – optimal PE as per the sawtooth hypothesis) and log(Clutch size) of egg masses; the black line is a fit from a Phylogenetic Least Squares (PGLS) model (F_1,76_ = 7.99, p = 0.0060, adjusted R^2^ = 0.08). In both panels, each point color represents a different pentatomid species (3-5 observations per species). (C) The protection differential (PD), circularity, and aggregation index of egg masses with clutch sizes that conform to neither hypothesis (n=32), the ovariole multiple hypothesis only (n=12), the sawtooth hypothesis only (n=13), or both hypotheses (n=21).

The shapes of egg masses with clutch sizes predicted by the sawtooth and ovariole multiple hypotheses were not consistent with more protective geometry. Protection efficiency (F_3,74_ = 5.3, p = 0.0023) and aggregation index (F_3,74_ = 4.25, p = 0.0079) varied among egg masses corresponding to each, both, and neither of the two hypotheses, and were highest on average for egg masses that conformed to neither hypothesis (Figure 3C). Circularity did not vary among egg masses in each category (F_3,74_ = 1.79, p = 0.16, Figure 3C).

Across all egg masses, more circular egg masses tended to have higher protection efficiency (*r* = 0.82, p < 0.0001, Figure S3). Aggregation index values correlated weakly with circularity (*r* = 0.21, p = 0.059) and no correlation between aggregation and protection efficiency was found (*r* = 0.08, p = 0.45, Figure S3). In other words, tightly packed (=high aggregation index) eggs were found in both circular egg masses with high protection efficiency as well as non-circular egg masses with low protection efficiency.

#### Intraspecific variation in clutch shape

Stink bug species differed in variation in clutch shapes, and thus protection efficiency. Species that laid smaller clutches had higher variation in protection efficiency with fewer egg masses conforming to the sawtooth hypothesis’s predictions; species with larger egg masses had protective geometry that was less variable and closer to optima predicted by the sawtooth hypothesis (Figure 4). Larger clutches tended to have better protective geometry (i.e., a greater protection differential) in *N. viridula* (*r* = 0.40, p = 0.0044), but not in *H. halys* (*r* = 0.20, p = 0.34), *E. heros* (*r* = -0.26, p = 0.20), or *C. impicticornis* (*r* = -0.25, p = 0.31). There was actually a slight negative correlation between protection differential and clutch size for *C. simplex* (*r* = -0.48, p = 0.00047); that is, larger clutches tended to have increasingly suboptimal protective geometry (albeit over a relatively small range of clutch sizes; Figure 4).

**Figure 4.**
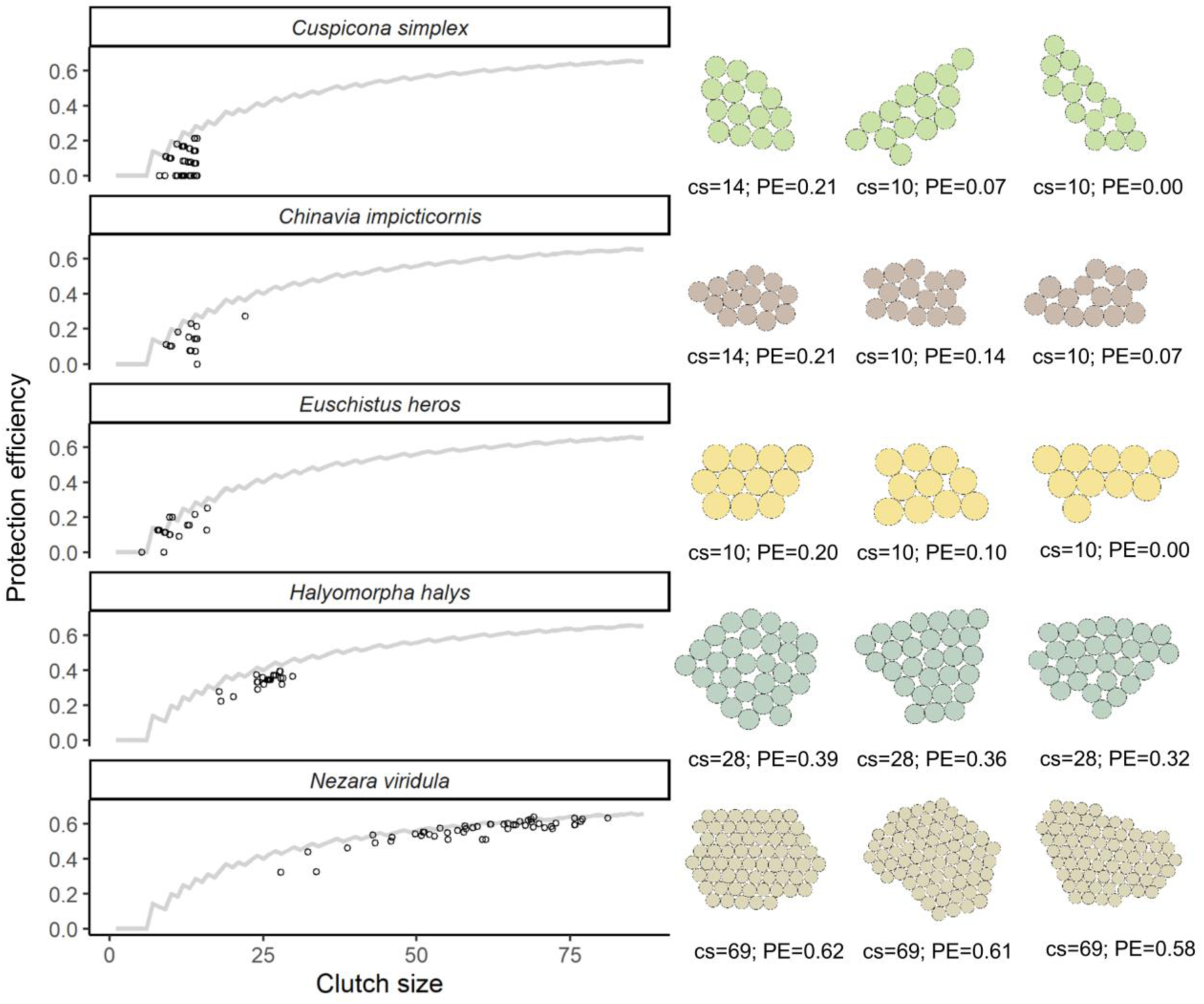
Examples of intraspecific variation in clutch shape in five species of pentatomid bugs. For each species, points show protection efficiency (PE) of individual egg masses; the grey lines show optimal PE for each clutch size predicted by the sawtooth hypothesis. Traced images of stink bug egg masses show examples of how egg masses with the same clutch size (cs) can have different shapes, and thus different PEs.

### Part 2: Clutch size variation

#### Clutch size analysis

Mean clutch sizes ranged across species from 6.8 eggs (*Cosmopepla lintneriana*) to 78.1 eggs (*N. viridula*) (Figure 5). Clutch sizes predicted by the sawtooth hypothesis were significantly over-represented in 16 species, significantly under-represented in 4 species, and not significantly different from the null scenario in the remaining 26 species (Figure 5C). Under-represented sawtooth-predicted clutch sizes occurred in species that laid many clutches that had less than 7 eggs – the lowest predicted adaptive peak in the sawtooth hypothesis – but whose clutch size distributions were right-skewed, increasing the distribution’s mean and thus the number of expected clutches with sawtooth-predicted clutch sizes, as well as in two species that often laid 28 eggs.

**Figure 5.**
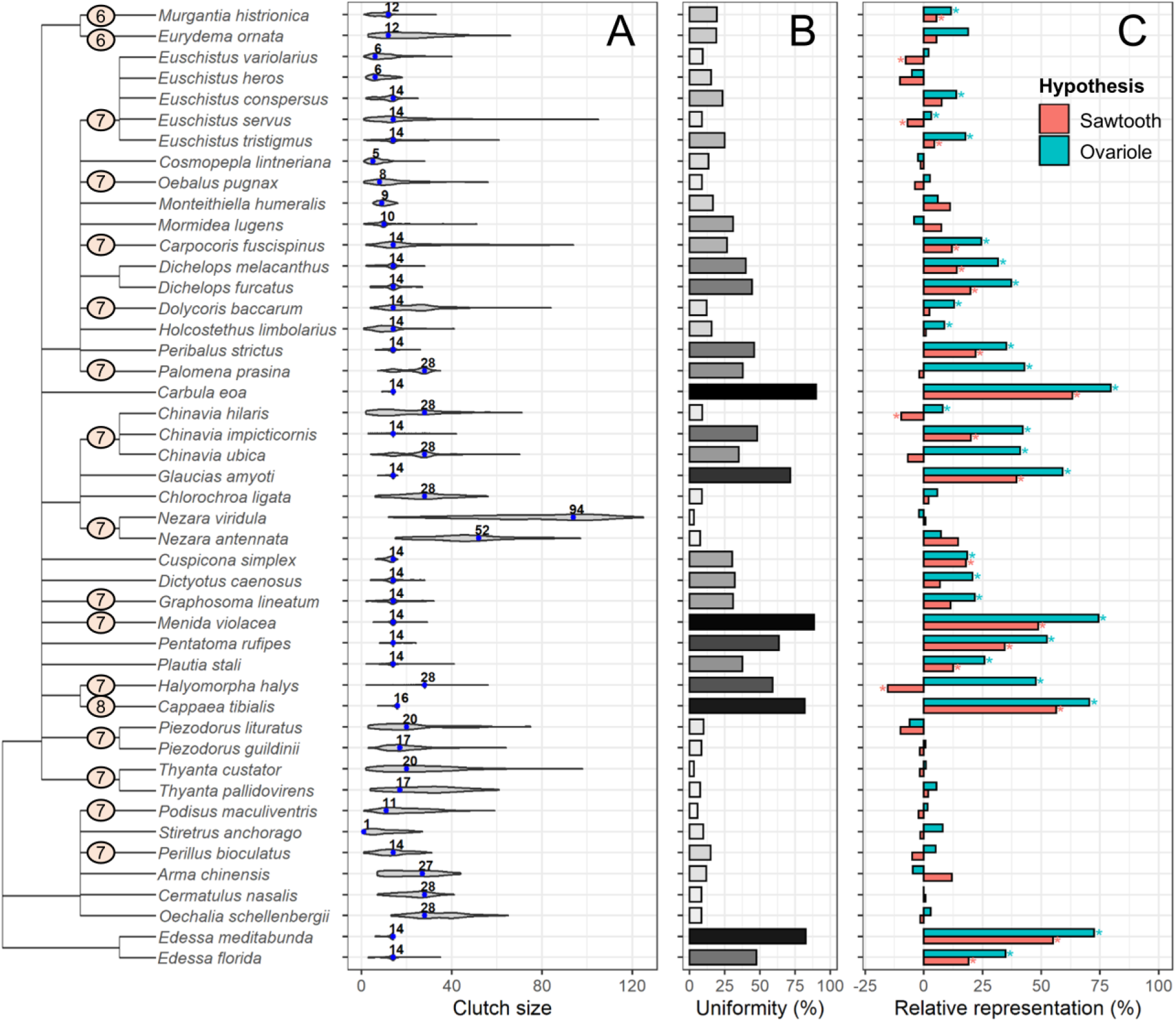
Clutch size variation within and among 46 species of stink bugs, arranged on a dendrogram representing taxonomic relationships. On the dendrogram, orange circles with numbers show the number of ovarioles per ovary for genera where this has been recorded. (A) Violin plots (normalized to have the same maximum width) showing distribution of clutch sizes; blue points with labels show the most common (modal) clutch size. (B) Percentage of egg masses that have the most common clutch size (darker colors are a visual aid indicating more uniform clutch sizes). (C) The relative representation (positive values = over-represented; negative values = under-represented) of clutches that had a number of eggs consistent with the sawtooth hypothesis versus the ovariole multiple hypothesis. In (C), an asterisk (*) indicates that the frequency of clutches matching a given hypothesis is significantly different than would be expected if clutch sizes were Poisson-distributed, as per 2×2 chi-squared tests of independence, controlled for multiple comparisons.

Clutch sizes predicted by the ovariole multiple hypothesis were significantly over- represented in 26 species, were not significantly under-represented in any species, and were not significantly different from the null scenario in 20 species. Among all 16 species for which both sawtooth and ovariole-multiple-predicted clutch sizes were over-represented, the representation of ovariole multiple-predicted clutch sizes was higher (mean difference = 15.2%; paired t-test: df = 15, t = 10.35, p < 0.0001; Figure 5C).

The degree of representation of clutch sizes predicted by the ovariole multiple and sawtooth hypotheses were highly correlated with clutch size uniformity (respectively, *r* = 0.96 and *r* = 0.84). That is, clutch sizes predicted by both hypotheses occurred more often in species that often laid the same-sized clutches. However, there were a few notable exceptions: species that often laid clutches of either 14 or 28 eggs (i.e., had bimodal clutch size distributions), such as *Chinavia hilaris* and *Dolycoris baccarum*, had low egg mass uniformity but strong over-representation of numbers predicted by the ovariole multiple hypothesis.

#### Clutch uniformity analysis

There was considerable variation in the uniformity of clutches sizes laid by the 46 stink bug species (Figure 5A). The most common modal clutch size was 14 eggs (observed in 21 species). The next most common modal clutch size was 28 eggs (seven species).

The uniformity of egg masses (the percentage of egg masses that conformed to the modal clutch size for a given species) varied from 3.0 to 89.8% (Figure 5B). Modal clutch sizes of 14 and 28 were the most common both for species with high clutch size uniformity and low clutch size uniformity (Figure 5B).

When accounting for shared ancestry, more uniform clutch sizes were most likely to occur at intermediate clutch sizes (contrary to the sawtooth prediction of lower clutch sizes) (Figure 6). High uniformity at intermediate clutch sizes was due to clutch sizes of 12, 14 and 28 having the highest uniformity (see above). Clutch size uniformity above 25% never occurred for species with average clutch sizes of below 9 eggs or of above 26 eggs (Figure 6).

**Figure 6.**
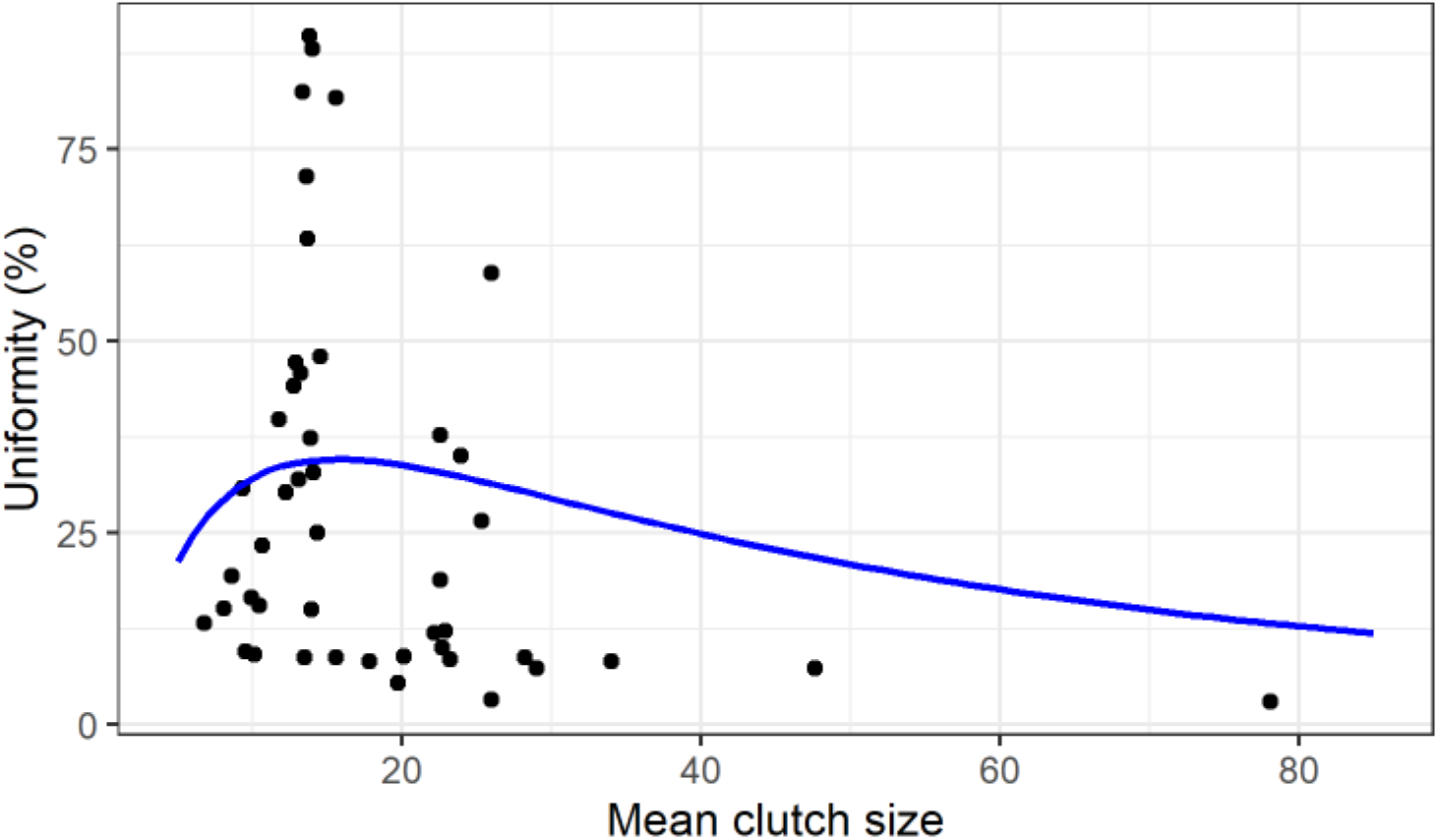
The relationship between mean clutch size and the uniformity of clutch size (% of egg masses conforming to the modal clutch size) in 46 species of stink bugs. The line is back-transformed from fitting log-transformed uniformity against mean clutch size measurements using a phylogenetic least-squares model including a polynomial term: log(uniformity) = -2.45 ×log(mean clutch size) – 1.81 × log(mean clutch size)^2^ + 3.08 (F_2,43_ = 7.59, p = 0.0015, adjusted R^2^ = 0.23).

#### Taxonomic patterns of clutch size and uniformity

There was a strong phylogenetic signal (based on taxonomic relationships) for mean clutch size (λ = 0.97, p = 0.0052), clutch size uniformity (λ = 1.11, p = 0.00037), and relative representation of ovariole multiples-numbered clutch sizes (λ = 1.06, p = 0.0014), but not for the relative representation of sawtooth-numbered clutch sizes (λ = 0.86, p = 0.087) (Figure 5).

Species within eighteen genera in 11 tribes of phytophagous stink bugs in the subfamilies Pentatominae and Edessinae laid egg masses with relatively high uniformity (>25%; mean ± SD: 49.9 ± 21.3%). Predatory stink bugs in the subfamily Asopinae (e.g., *P. maculiventris*) all tended to lay less uniform egg masses (5.4–15%; mean ± SD = 9.8 ± 3.3%). Low clutch size uniformity was also present in some pentatomine genera such as *Thyanta* and *Nezara*. There was, in some cases, notable variation in these parameters even within genera. One genus, *Euschistus*, included two species with high-uniformity clutch sizes with over-representation of ovariole multiples, as well as two species with low-uniformity clutch sizes (Figure 5).

## Discussion

For oviparous animals, the number of eggs laid per reproductive bout (clutch size) has been the subject of extensive theoretical and empirical work, putting forward and testing hypotheses about how optimal and realized clutch sizes have evolved (Godfray et al. 1991; Wilson and Lessells 1994). In general, mothers are expected to produce clutches with a number and size of eggs that maximize successful offspring production over their lifetimes, but with clutch size optima affected by environmental stochasticity and constrained by resource availability and life history tradeoffs (e.g. Parker and Courtney 1984; Ives 1989; Roitberg et al. 1999; Olofsson et al. 2009). Another type of constraint on the production of optimally sized clutches could be the reproductive anatomy of the mother, which is known to influence total reproductive output (Sarikaya et al. 2019; Church et al. 2021), but its association with clutch size has not been thoroughly examined. We tested two hypotheses to explain why some species of stink bugs often lay clutches containing the same number of eggs: one related to how egg mass geometry may protect against environmental challenges and the other related to reproductive anatomy.

The “sawtooth hypothesis”, predicting that stink bug clutch sizes have been selected to converge on numbers of eggs that offer the best protective geometry, is an appealing hypothesis to link animal clutch size to clutch geometry. This hypothesis was inspired by a number of past studies showing strong mortality differences between inner and outer eggs in egg masses of true bugs (McLain and Mallard 1991; Mappes et al. 1997; Kudo 2001; Kudo 2006). However, our analysis does not support the sawtooth hypothesis, at least not for stink bugs. Especially for smaller clutch sizes, stink bugs often do not lay egg clusters in a shape that would protect the maximum proportion of eggs, even when laying a clutch size (e.g., 14) that could be arranged in a shape predicted to lie on an ‘adaptive peak’. In fact, our data show that several species often lay clutches of eggs that do not protect any eggs in the interior (Figures 3, 4). However, we did find that the relationship between predicted protection efficiency and clutch size had the same general form as predicted by the sawtooth hypothesis, with real clutch geometries protecting increasingly optimal numbers of eggs in clutch interiors (i.e., becoming more hexagonal) as clutch size increased; this pattern was present interspecifically (Figure 3B) as well as intraspecifically in one species (Figure 4). Thus, contrary to our initial prediction that selection for optimal clutch geometry would most strongly operate at smaller clutch sizes (because it would provide the greatest *proportional* increase in protection efficiency), our results better support the hypothesis that selection for protective geometry may be stronger at higher clutch sizes because the opportunity for increasing the *number* of offspring protected at the interior is greater. For these very large clutches, the marginal gain or loss in protective geometry from adding or subtracting an egg would be minimal, and so selection for uniformity would be unlikely. Based on our results, we make the following prediction: for animals that lay their eggs in clusters, all else being equal, larger clutches should be more hexagonal but not more uniform in egg number. This general phenomenon of larger aggregations being more hexagonal may also apply to aggregations of a variety of other animals for which there is higher risk of mortality on the edges of aggregations (Hamilton 1971; McClain and Mallard 2019).

We found strong support for the idea that the number of ovarioles in stink bug ovaries, usually 7, might influence the most common sizes of clutches, as many species were found to usually lay clutches of 14 or 28 eggs. In further support of the ovariole multiple hypothesis, in two species (*M. histrionica* and *E. ornatum*) known to belong to a taxonomic group in which species have a total of 6 ovarioles in each of their two ovaries, the most common clutch size was 12 and, in *C. tibialis*, which has eight ovarioles per ovary, the most common clutch size was 16. The relationship between clutch size uniformity and ovariole number has previously been hypothesized in individual species or closely related groups of stink bugs (Kiritani and Hokyo 1965; Panizzi and Herzog 1984; Matesco et al. 2008; Matesco et al. 2009) but our study shows that this is a broad-scale trend, the prevalence of which appears to cluster, to some degree, with taxonomic relatedness. Our results, however, also show that high clutch size uniformity is not inevitable when ovariole number is essentially fixed: for 20 out of 46 species in our dataset, ovariole number multiples were not over-represented in the clutch size distributions. These findings build on recent studies of insects (Sarikaya et al. 2019; Church et al. 2021) recognizing the broad-scale importance of ovariole number as a key aspect of reproductive anatomy underlying variation in animal reproductive strategies. In Hawaiian *Drosophila*, variation in ovariole number has rapidly evolved during an adaptive radiation associated with different ecological niches (oviposition substrates and levels of specialization) and is positively associated with potential fecundity (Sarikaya et al. 2019). However, high inter-taxa variation in ovariole number is by no means the rule; it has recently become apparent that many groups of insects have evolved fixed or nearly-fixed ovariole numbers (Church et al. 2021). Our study shows that even within a group of insects that has strong consistency in ovariole number, a wide variety of clutch sizes and degrees of clutch size uniformity has evolved.

The likely mechanism underlying variation in clutch size for species with the same ovariole number is the ability to retain different numbers of mature oocytes before laying; for example, species or individuals that often lay 14 and 28 eggs mature one or two oocytes per ovariole (= 7 or 14 per ovary) before laying, respectively (Matesco et al. 2009). The fact that the observed degree of clutch size uniformity varies considerably among the species in our dataset, to the point where some species have highly variable clutch sizes that are not multiples of ovariole number implies that oocyte retention may be an extremely plastic trait for some species. Therefore, stabilizing selection in response to environmental challenges might not be the major influence on patterns of clutch size uniformity (as inferred by the sawtooth hypothesis). Uniformity in some species may in fact be largely due to ovariole number, with selection in other species operating to promote plastic clutch size strategies that result in decreased uniformity. What life history and ecological factors could be associated with a transition away from uniformity and towards plasticity in clutch size? Although a detailed analysis of the ecological correlates of clutch size uniformity is outside the scope of this study, the taxonomic patterns we observed allow us to put forward some hypotheses to guide future research.

Higher plasticity in reproductive strategies are expected to evolve under conditions of higher environmental variability. One possibility is that for insects that lay their eggs in clusters, the ability to partially ‘break free’ of constraints imposed by reproductive anatomy and evolve the capacity to vary clutch size is favoured when the quantity or quality of food resources for oogenesis and/or for offspring hatching from the clutch is highly variable and/or unpredictable over time. For example, predatory stink bugs in the subfamily Asopinae have highly variable clutch sizes, and multiples of ovariole number are not obviously over-represented. The size of prey items consumed by predatory stink bugs in nature can be highly variable (P.K.A, personal observations), and there is evidence that intervals between prey captures can also be variable (Legaspi et al. 1996). Being able to adjust clutch sizes according to the amount of food available to support oogenesis would allow predators to produce clutch sizes that correspond with their nutritional status or the expected resource availability for their offspring (De Clercq and Degheele 1992; Legaspi and O’Neil 1993; Ware et al. 2008). Accordingly, numerous studies have shown that mean clutch size can vary considerably under different environmental conditions (e.g., food limitation) in predatory stink bugs (e.g., De Clercq and Degheele 1997; Adams 2000; Lemos et al. 2006; Torres-Campos et al. 2016). Highly uniform clutch sizes are more common in the subfamily Pentatominae, whose members are mostly phytophagous and for which food resource availability would be expected to be more predictable. However, some phytophagous genera (e.g., *Cosmopepla*, *Thyanta*) produced variably-sized clutches that do not fit this general pattern. One possibility is that these species have a slow or uncertain rate of oogenesis (e.g., due to variation in host plant nutritional quality) relative to their risk of mortality, and thus may be selected to lay clutches before they can mature a complete complement of oocytes. Ageing, which can influence the functional number of ovarioles (Prathibha and Krishna 2010), may also be an important factor contributing to reducing clutch size uniformity, and could have contributed to some of the variability in our clutch size dataset as well. Interrelationships between resource intake, mortality risk, age, and clutch size variation is a promising area for future studies in this group of insects as they exhibit such a wide variety of life-history variation.

What influences the wide variation in observed clutch size geometry? Although we found that larger clutches converged towards being almost hexagonal, producing large clutch sizes, that protect large numbers of interior eggs, might not be possible for many species due to constraints such as body size; some of the species in our dataset that laid the largest clutches are also among those with the largest body sizes (e.g., *N. viridula*, *C. ligata*, *C. hilaris*, Paiero et al. 2013). Smaller species of stink bugs often laid clutches of less than 14 eggs, which were sometimes laid in linear 2-by-n rows (e.g., *M. histrionica*, *Oebalus pugnax*, *Stiretrus anchorago*, *Piezodorus guildinii*). Eggs within these clutches are as aggregated as eggs within larger ones, but their arrangements do not provide protection to inner eggs. Other species, such as predators in the genus *Podisus*, often lay relatively small, loosely clustered (non-aggregated) eggs with highly suboptimal protective geometry. There is some evidence that in some of these species with clutch sizes too small to protect a large number of eggs, other strategies have evolved to protect eggs from predation and/or parasitism, such as camouflage, occupying enemy-reduced microhabitats (Torres-Campos et al. 2016) or sequestering toxic chemical compounds from host plants (Guerra-Grenier et al. 2021). In the latter case, there is preliminary evidence that egg patterning may act as an aposematic signal to predators; if so, laying eggs in 2-by-*n* lines would maximize the strength of this signal. The shape and structure of oviposition substrates, as well as internal factors (e.g., age, nutritional stress, mating status), are likely to be further important constraining factors for clutch shape and size (Silva and Panizzi 2008; Figure 2G, K). Future studies would need to establish relationships between clutch size (and its uniformity) and fitness to determine to what extent observed clutch size patterns reflect historical processes of selection and adaptation versus environmental constraints.

Our results highlight an interesting link between reproductive anatomy and ecology: at least in some species of insects, knowledge of ovariole number can help to predict the most common clutch sizes they will lay, and vice versa. This correspondence may be most likely to exist in species that coordinate their oocyte maturation and lay eggs in aggregated clutches with highly uniform sizes. Our study suggests that such species may typically lay intermediate-sized clutches. Such patterns may be a good preliminary indication that ovariole number may be influencing the sizes of the most commonly observed clutches. Examining these relationships in other groups of animals with variable and invariant reproductive anatomy, and where there are different selective pressures on clutch size geometry, will undoubtedly continue to reveal interplay between development and ecology and continue to advance research on the ecology and evolution of insect eggs (Donoughe 2021).

## Acknowledgements

We thank Dr. Toshiharu Mita (Kyushu University) for providing data from Japan, and Talita Roell for help with literature searches. I.C.W.H. and P.K.A. acknowledge support from the Israel Institute for Advanced Studies for the research group programme ‘Mathematical modeling of biological control interactions to support agriculture and conservation’. We thank the many people who helped with insect culture maintenance and field collections that generated clutch size data for this manuscript. We also thank Sam Church (Harvard University) for helpful discussions regarding an earlier version of this manuscript.

## Supplementary Material

**Table S1.**
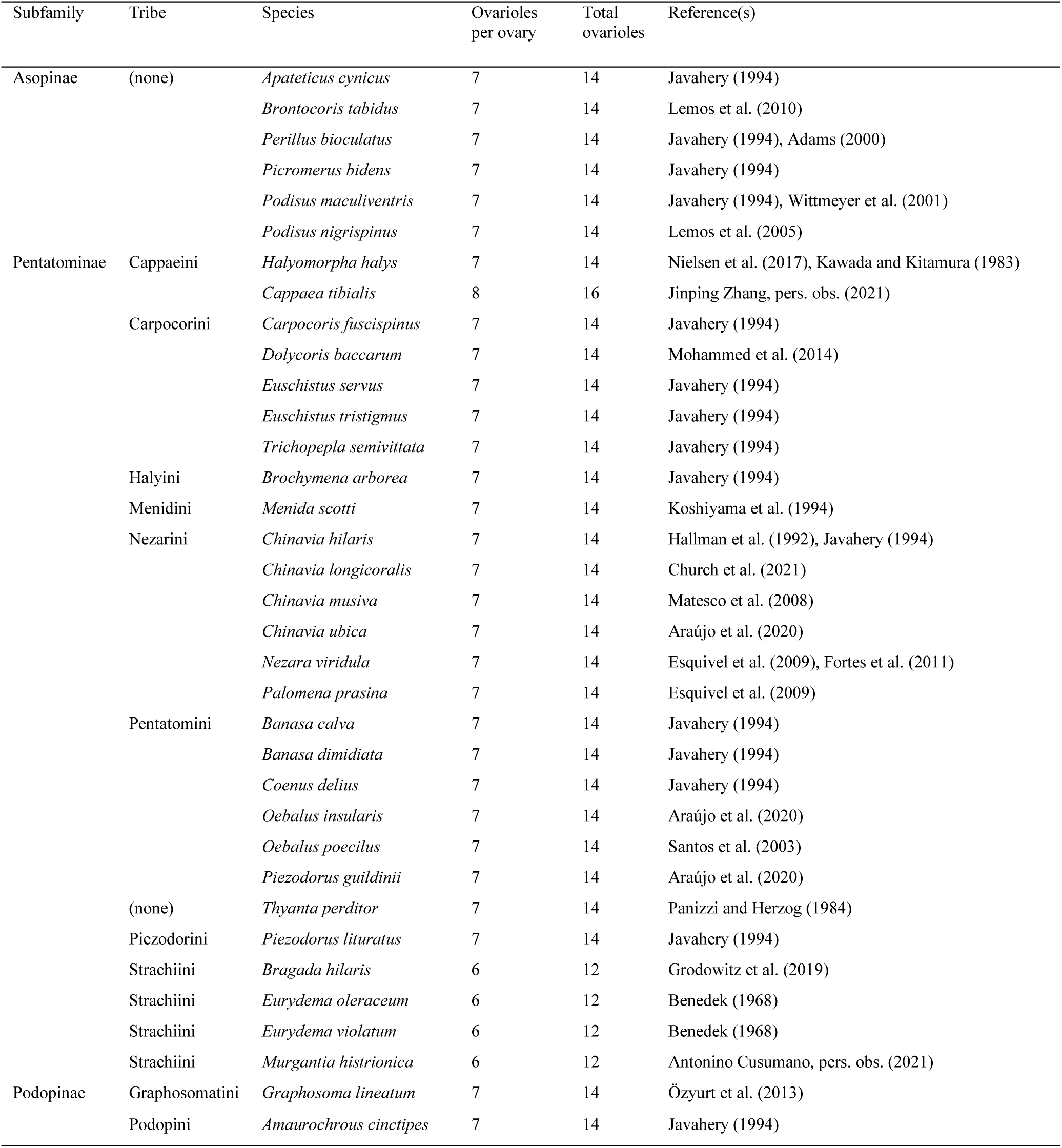
The number of ovarioles per ovary and the total number of ovarioles of 35 species of pentatomid bugs.

**Figure S1.**
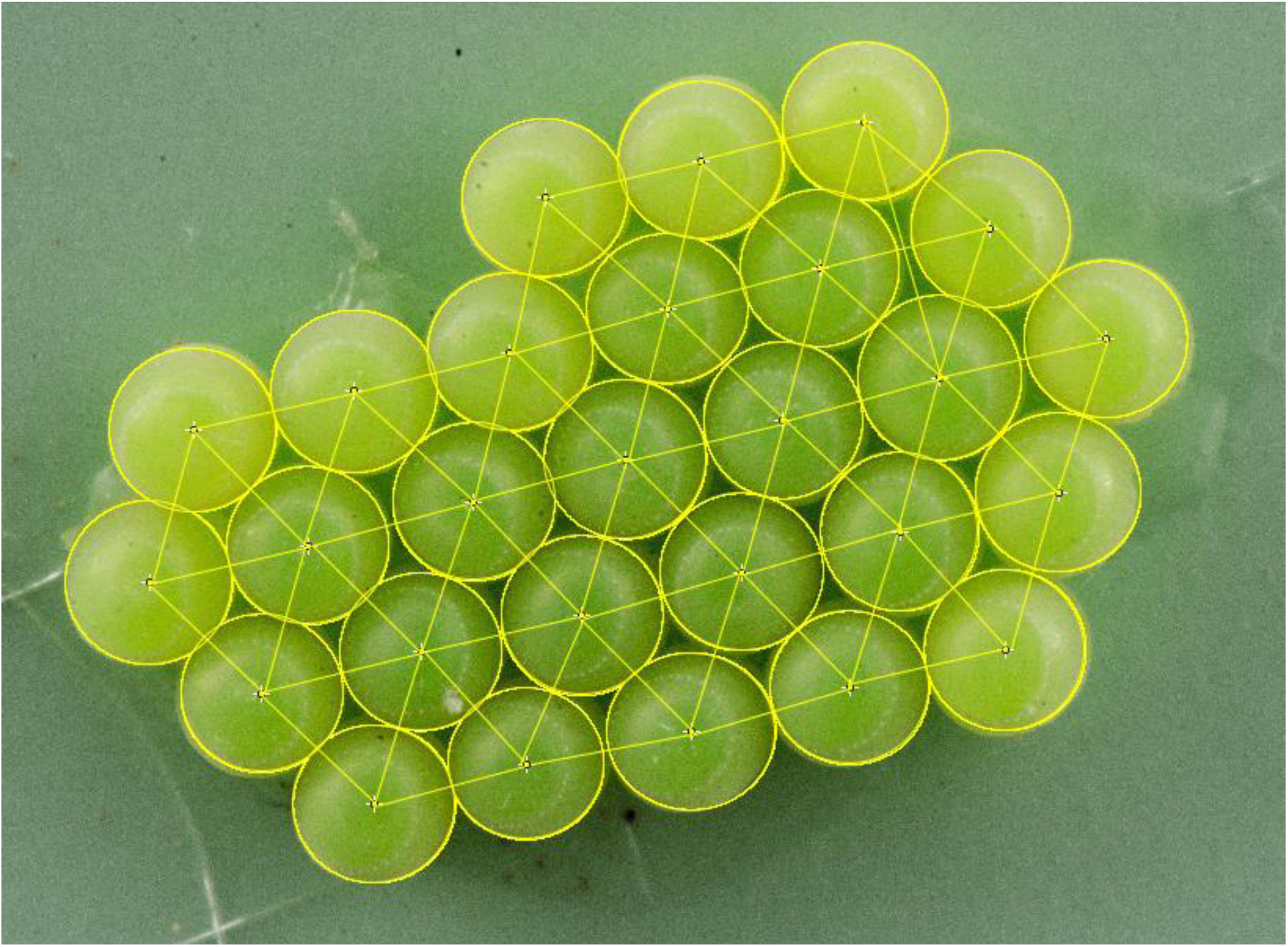
An example of regions of interest (ROIs) drawn on a *Palomena prasina* egg mass in ImageJ to measure clutch parameters associated with geometry.

**Figure S2.**
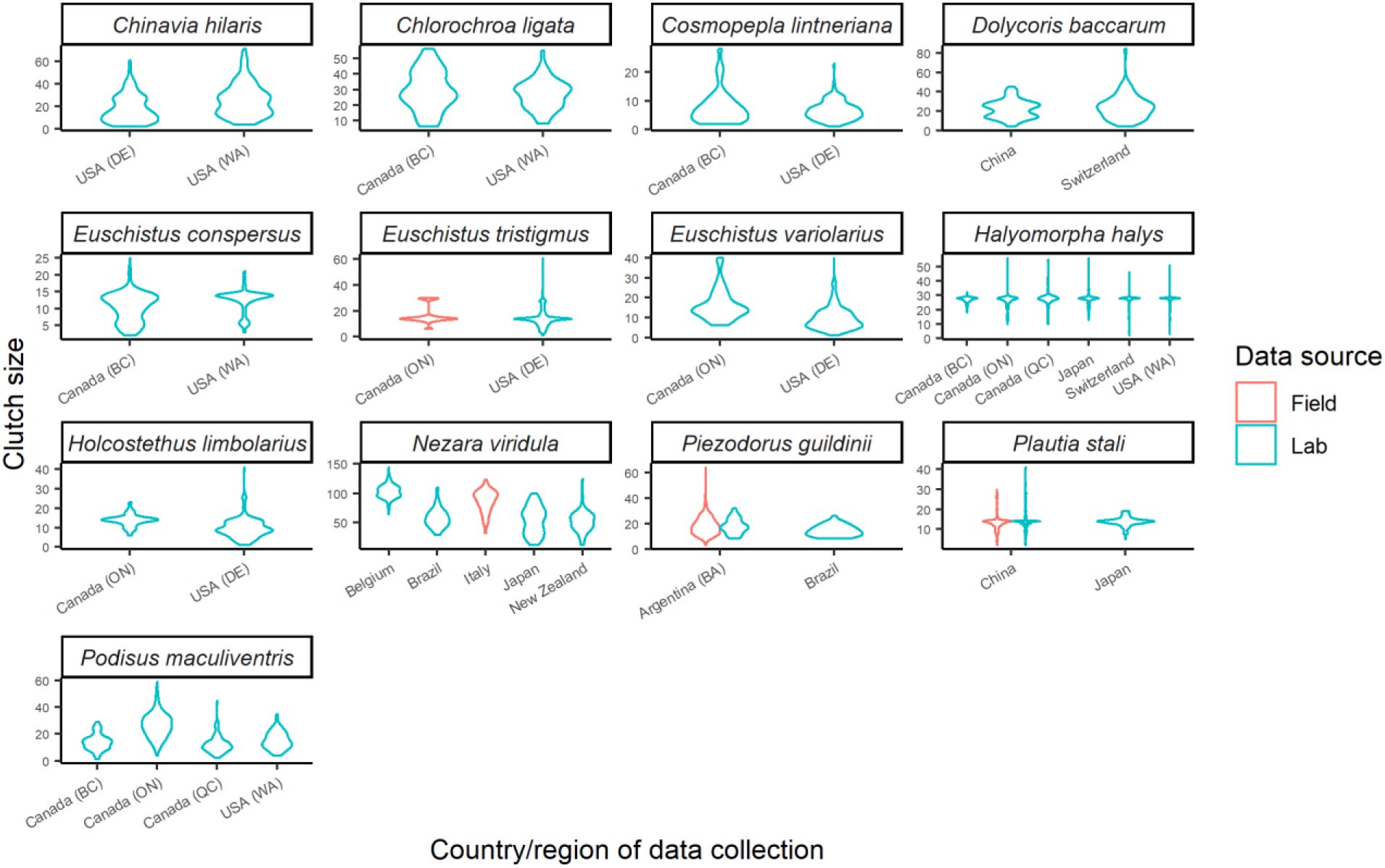
Consistency of variation in observed clutch sizes among datasets for the same Pentatomid species in datasets from the laboratory and the field, and from different research groups. Only species for which there were at least two independent datasets with at least n=30 egg masses per dataset are shown. Violin plots are normalized to have the same maximum width and each y-axis is different. For the USA and Canada from which there were regional datasets, results are shown according to state or province, respectively.

**Figure S3.**
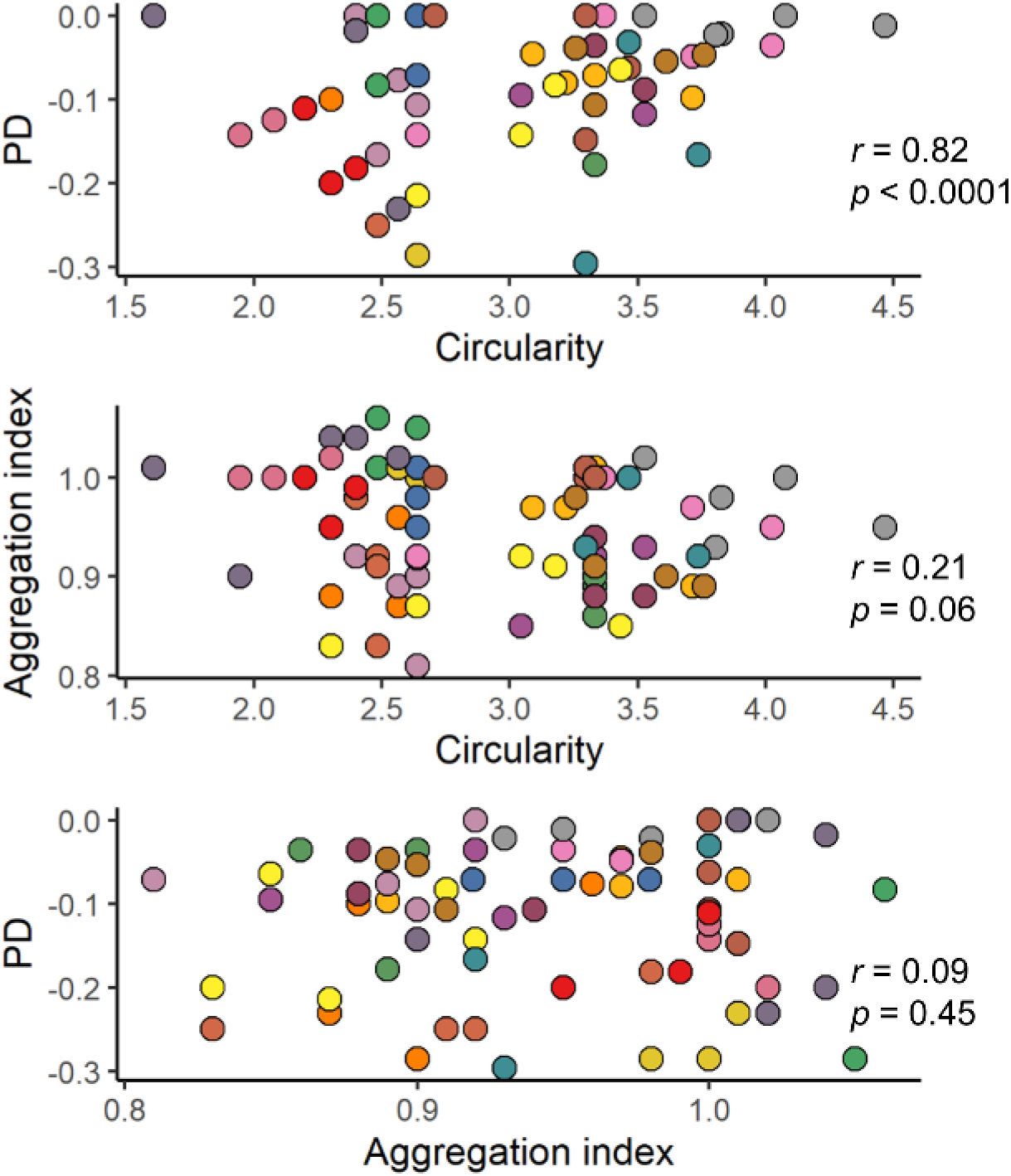
Correlations among protection differential (PD), aggregation index, and circularity measured from 78 pentatomid egg masses (belonging to 19 species) in relation to the sawtooth and ovariole multiple hypotheses. Each point color represents a different stink bug species. Statistics are from Pearson’s correlations.

## Footnotes

1 For an engaging explanation of why “hexagons are the bestagons”: https://www.youtube.com/watch?v=thOifuHs6eY.

